# Causal modeling dissects tumour-microenvironment interactions in breast cancer

**DOI:** 10.1101/144832

**Authors:** Leon Chlon, Florian Markowetz

## Abstract

Elucidating interactions between cancer cells and their microenvironment is a key goal of cancer research with implications for understanding cancer evolution and improving immunotherapy. Previous studies used association-based approaches to infer relationships in transcriptomic data, but could not infer the direction of interaction. Here we present a causal modeling approach that infers directed interactions between signaling pathway activity and immune activity by anchoring the analysis on somatic genomic changes. Our approach integrates copy number profiles, transcriptomic data, image data and a protein-protein interaction network to infer directed relationships. As a result, we propose 11 novel genomic drivers of T cell phenotypes in the breast cancer tumour microenvironment and validate them in independent cohorts and orthogonal data types. Our framework is flexible and provides a generally applicable way to extend association-based analysis in other cancer types and to other data and clinical parameters.

## Background

Solid tumours like breast cancers are complex tissues consisting of a cell-autonomous compartment of cancer cells accumulating somatic changes and undergoing clonal evolution [1] and a non-cell-autonomous compartment containing lymphocytes, fibroblasts and other cell types forming the tumour microenvironment [2]. Both compartments are known to influence each other [3] but the details of how they communicate are still poorly understood. For example, it is unclear how breast cancer cells hijack signaling pathways and regulatory mechanisms to influence the recruitment and activity of tumour infiltrating lymphocytes (TILS) [4, 5].

Previous studies used association-based methods to characterise tumour-micro-environment interactions within the bulk [6, 7] and micro-dissected tumour transcriptomes [8]. For example, Ali *et al* [7] showed that patterns of immune infiltration varied between molecular subtypes of breast cancer in bulk tumour transcriptomes; and Oh *et al* [8] derived stromal-epithelial co-expression networks from micro-dissected tumour data to investigate crosstalk within the tumour microenvironment.

Data from transcriptomic studies are readily available, and association-based analyses have provided profound insights into the general structure of co-expression patterns within the data. However, association-based methods have several limitations. First of all, when analysing bulk data they are limited by the convolution of gene expression patterns arising from both the cell autonomous and non-autonomous compartments [9]. Second, associations alone can not distinguish whether a change in gene expression is caused by a change in immune activity or is a response to it [10].

To overcome these limitations, we propose a statistical approach integrating data on somatic genomic changes, signaling pathway activity and immune activity in the tumour. Our approach assigns a direction to an association between signaling pathway activity and immune activity by anchoring the analysis on an underlying genomic event. We exemplify this general idea here by using copy number alterations (CNA) to measure genomic events, transcription factor (TF) activity inferred from transcriptional profiles to measure signaling activity, and expression of marker genes as well as imaging data to measure immune activity. Based on these data, our approach uses Bayesian networks and likelihood approaches to choose the best fitting model (similar to [10]), and reduces the search space by using protein-interaction networks to limit the number of potential models.

For discovery, we apply our approach to 1,980 breast tumour samples with paired genomic-transcriptomic data [11] to generate candidate models for transcriptomic phenotypes. For validation, we test these models in an independent genomic-transcriptomic dataset comprising 1,154 samples [12]. For further validation on an orthogonal measure of immune activity, we use data on lymphocyte infiltration derived from over 550 Haematoxylin & Eosin (H&E) stained whole tumour slides [13]. As a result, we find 12 models that robustly validate and propose 11 novel genomic drivers of T cell phenotypes in the breast cancer tumour microenvironment.

## Results

We developed a framework to formalise causal relationships between somatic genomic events, signaling pathway activity and immune activity in the tumour. Our approach is implemented in the statistical environment R [14] and all code to reproduce the results presented here is available as part of a annotated document in the supplementary information.

To measure genomic events we used the copy number profiles of 19,702 genes, as provided by the METABRIC [11] and TCGA projects [12]. To measure signaling pathway activity, we focused on 788 experimentally verified TFs [15]and used the paired transcriptional profiles from the same resources to apply VIPER [16], a method for network based prediction of transcription factor activity.

To measure immune activity, we used two orthogonal approaches: the first approach uses the mean expression of marker genes to define a cytolytic score (CS) [17] and a novel T-cell score (TCS). While the CS trait is a measure of lymphocyte activity, the TCS measures the degree to which they are represented in the tissue. We validated TCS on paired gene expression and flow cytometry blood sample data [9] and found that of the 9 leukocytes subsets tested, CD8+ T-cells demonstrated the only significant positive correlation with the TCS (*ρ* = 0.675, *P* = 0.001), while the remaining leukocytes show either negative or no significant correlation (see Additional file 1). The second approach uses paired H&E images from the METABRIC cohort to measure the absolute number of lymphocytes and their density per tumour [13]. The details of how we derived these measurements are given in the Materials and Methods section.

### A multi-step causal inference approach to assign directionality to signaling-immune associations

Our framework to assign directionality to signaling-immune associations is motivated by an established approach to order gene expression traits relative to one another and relative to other complex traits [10]. The key idea is to anchor the analysis on genomic variation and to systematically test whether DNA changes that lead to changes in signaling and immune activity support a causative, reactive or independent model of the interaction between signaling and immune activity.

We formalise different causal relationships in three different types of graphical models (Fig. 1A). Model 1 (*M*_1_: the causative model) represents a case in which a genomic event changes immune activity by perturbing signaling activity. Model 2 (*M*_2_: the reactive model) represents a case in which a genomic event leads to a change in immune activity, which then in turn changes signaling activity. Model 3 (M3: the independence model) represents a case in which the genomic event influences immune activity and signaling activity independently of each other. We used standard assumptions of causal inference [18] to derive likelihood functions for each of the three models (see Materials and Methods). For each triplet consisting of one genomic locus, one transcription factor and one immune score, we maximise the likelihood function of the three models over their parameters, and finally choose the model with the smallest Akaike Information Criterion [19], a model selection criterion balancing goodness-of-fit with model complexity.

**Figure 1.**
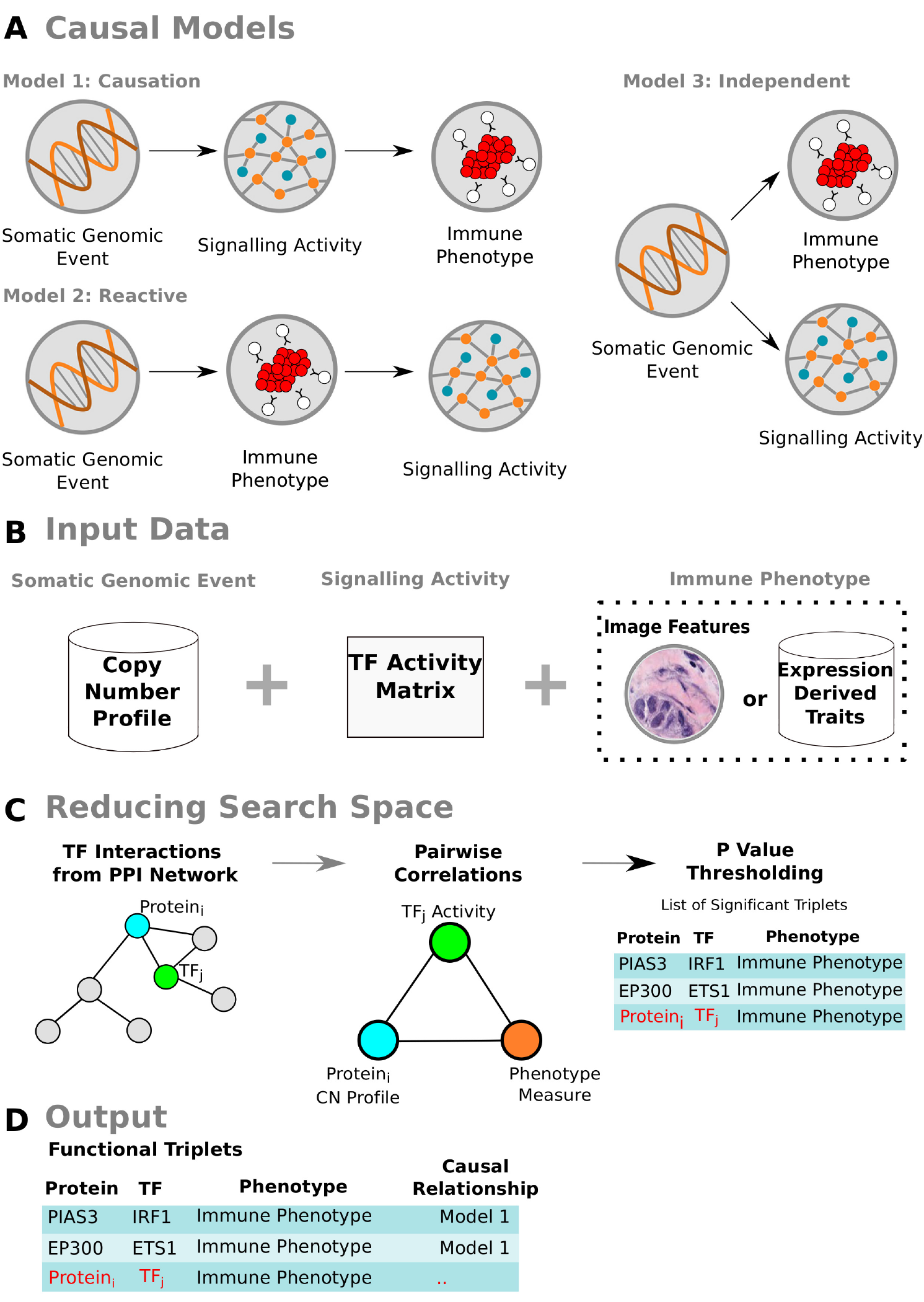
Description of CMIF. **A** Directed Acyclic Graphs (DAGs) representing each respective model evaluated during the analysis. Model M1 describes a causal relationship in which the DNA aberration event acts on the immune trait through perturbation of the underlying transcription factor network. Model M2 described as a reactive relationship with respect to the transcription factor activity, causally modulated by the immune trait. Model M3 describes an independence relationship, in which the DNA aberration acts upon each of the traits independently. **B** The inputs for CMIF are a matrix of TF activities per sample as measured by VIPER, continuous intensity profiles for the copy number calls and phenotype data that can be either image features or features derived from gene expression data. **C** The CMIF extracts all experimentally verified interactions between a given TF and other intracellular proteins from a protein interaction network. Three pairwise correlations are then computed, the first between the copy number profile of the gene coding for the protein and the given TF’s activity, second between the copy number profile and the phenotype, and finally, between the TF activity and the phenotype. If all associations are significant at a user-defined threshold, the protein-TF-phenotype triplet is admitted to the next step of the analysis. Pre-defined likelihood functions for each model in **A** are maximised over their parameters using an algorithm for maximum likelihood estimation, with the model most likely to be supported by the data determined by the model with the smallest Akaike Information Criterion (AIC). **D** The output of CMIF is a table of functional triplets with their corresponding model classification.

To limit the search space and reduce the number of triplets to test, we developed a multi-step causal model inference framework (CMIF) approach (Fig. 1). In a first step, CMIF filters only selects genes with experimentally verified protein-protein interaction (PPI) with a TF of interest (Fig. 1C). There are 19, 702 × 788 = 15, 525, 176 pairwise associations between copy number profiles and TF activities, and filtering them according to the StringDB database [20] results in just 2,333 potential models. This filtering substantially reduces the search space and enriches for biologically relevant drivers in groups of correlated genes that are jointly amplified or deleted.

In a second step, undirected skeletal association graphs are constructed for CNA events underlying both the TF activity and immune phenotype by computing pairwise correlation coefficients between the variables. Benjamini-Hochberg [21] p-value correction is applied and only skeletons where all pairwise associations are significant are passed to the final step (Fig. 1C). Finally, the likelihood function of each model consistent with these skeletons is maximised over its parameters and the model with the smallest Akaike Information Criterion is chosen. After filtering and model selection, CMIF provides as output the model of the causal relationship between the variables (Fig. 1D).

### Evaluating CMIF with CS/TCS immune metrics

In the first analysis we used the TCS and CS metrics derived from gene expression data as measurements of immune activity. Applying CMIF to the TCS/CS metrics, TF activities and copy number profiles resulted in 111 unique TFs that were significantly associated with either the TCS or CS measurements, and whose underlying CNA modulator correlates significantly with the immune phenotype. CNA events at loci corresponding to 176 genes significantly correlated with both the TF activity and either immune phenotype, resulting in 475 undirected skeletal graphs. For each graph, we fit the likelihood models M1, M2 and M3 to the respective copy number profiles, TF activity and TCS/CS trait measurements (see Methods). 344 triplets (72%) were best represented by the causal model (M1) whereas the reactive model (M2) was the best in the remaining 131 (28%) cases. No M3 models were supported by the data, which can be explained by the degree to which the association and PPI filtering step of the method (Fig. 1C) identifies strictly causal or strictly reactive mechanisms.

#### Validation in independent cohort

We validated these results in the independent TCGA cohort. Of the candidate triplets, 194 (54.6%) M1 and 24 (18.3%) M2 models validated in the TCGA cohort using CMIF (Fig. 2A). The higher validation percentage of M1 models over M2 models in TCGA data indicates that causal drivers of T cell infiltration are more robust and thus more frequently recapitulated in breast cancer populations. The higher prevalence of M1 over M2 models might be explained by cancer cells being immunoedited [22], a process in which somatic mutations break downstream pathways associated with a normal immune response. Over time, this would enable the tumour to exert more control over the immune system than vice-versa.

**Figure 2.**
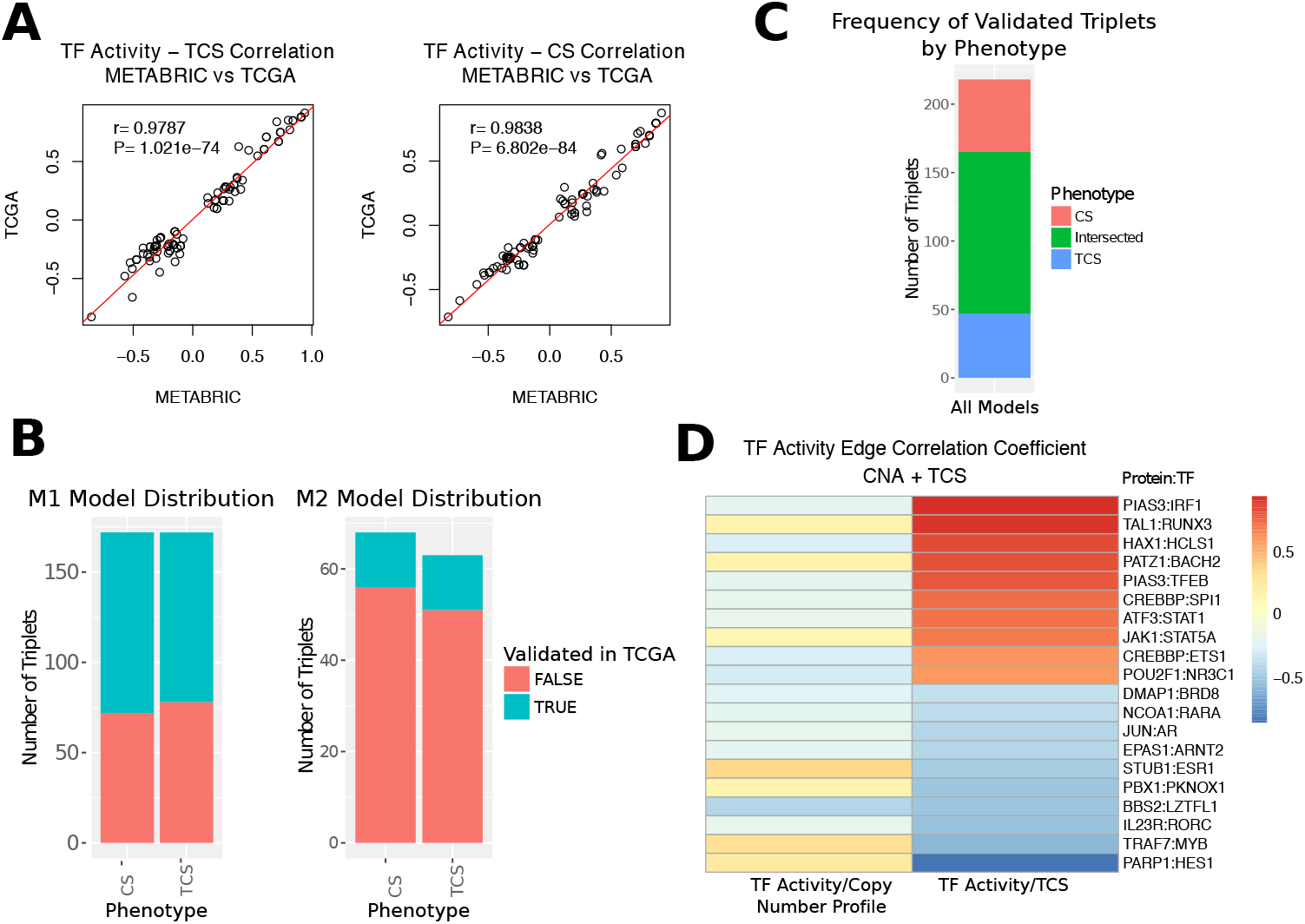
Model validation in TCGA. **A** Scatter plots with fitted regression lines illustrating strong concordance between METABRIC and TCGA when transcription factor activity significantly explains the variance in the TCS and CS immune traits. **B** The proportion of predicted relationships in METABRIC that validate in TCGA as stratified by model type and lymphocyte trait. **C** Stacked barchart illustrating the frequency and overlap of models between the different immune traits. **D** Top and bottom 10 TCGA-validated causal models as ranked by the proportion of TCS variance explained by the transcription factor activity. Y-axis indexing is organised as (Gene at locus of CNA event): (Transcription Factor). Heatmap columns illustrate Pearson’s correlation coefficient between the CNA signal and transcription factor activity, and transcription factor activity and TCS measurement (left to right).

The correlation between TF activity and the individual immune traits agreed well between METABRIC and TCGA (TCS: *ρ* = 0.98, *P* < 2.2 × 10^−16^; CS: *ρ* = 0.984, *P* < 2.2× 10^−16^) (Fig. 1B), highlighting robust co-dependence relationships between lymphocyte infiltration/activation and TF activity. Of the validated models, 118 were shared between the TCS and CS traits, with 47 unique to the TCS (165 total) and 53 unique to the CS (171 total). (Fig. 2C). This high degree of concordance is reassuring considering that many molecular pathways facilitating lymphocyte aggregation will also directly or indirectly influence the cytolytic activity and vice-versa.

#### Validation by literature

Many of the top predictions generated by CMIF are well supported by the literature. For example, when ranked by the strength of their correlation with the immune phenotype, CMIF analysis highlighted *I RF*1 as the strongest causal mediator of the TCS phenotype across both METABRIC and TCGA, whose TF activity is down-regulated by the amplification of *PIAS*3 (Fig. 2D). This is consistent with reports that *PIAS*3 induces transcriptional repression of IRF1 through binding to it as a SUMO-1 ligase [23]. Furthermore, *I RF*1 has been shown to play a crucial role in driving anti-tumour immune response [24] and thus this model’s categorisation as causal for TCS is well supported by the strong body of literature surrounding the relationship between the variables.

*RUN X*3, a well known tumour suppressor gene [25], was identified as the second strongest causal modulator of the TCS. *RUNX*3 activity has been found experimentally to mediate lymphocyte chemotaxis through the TGF-*B* pathway [26, 27]. The positive association found between *TAL1* and *RUNX*3 has been also been confirmed in studies demonstrating that the *RUNX* genes are direct targets of *TAL*1 [28]. Additionally, CMIF identified *ETS*1 as a causal mediator for the TCS, which is unsurprising given that its activity has been shown to regulate the transcription of chemokines and cytokines directly involved in lymphocyte migration [29].

#### Validation with image-derived features

Another way of validating the robustness of our model is testing how well it predicts lymphocyte infiltration in an orthogonal dataset. To facilitate this, we used the paired tumour whole tissue section slides stained with Haematoxylin & Eosin (H&E) provided by the METABRIC study [11], which provide an estimate of lymphocytic infiltration independent of the gene expression based estimates used above.

We evaluated the predictive utility of 165 validated TCS M1 models on an image cohort consisting of 534 samples. We used the results of previous image analyses to segment and quantify the absolute number of lymphocytes and the lymphocyte density [13], with further normalisation techniques applied in our study to generate traits from these features (See Methods).

We combined these image features, TF activities and copy number profiles in our CMIF approach and computed the overlap between the image-based causal models and those from the transcriptomic phenotype set. 18 (10.9%) of the initial predictors of the TCS were also predictive of the image features, with the majority (15/18) belonging to the lymphocyte density trait. The extent to which TF activity significantly correlated with both the TCS and the image lymphocyte density feature simultaneously was weaker (*ρ* = 0.45, *P* < 2.2 × 10^−16^) than that of the TCS between METABRIC and TCGA. This might be due to the fact that the transcriptomic T cell features are not necessarily perfect proxies to lymphocyte features extracted from images. For example, while the TCS makes measurements about T cells exclusively, the feature information in H&E images is not sufficiently descriptive to differentiate NK cells from T cells leading to a weaker correlation. Furthermore, there are confounding systematic errors that may have arisen during the segmentation and classification process used to generate the image features that render the correlation with the transcriptomic phenotypes weaker than expected.

For consistency, we only considered causal models that demonstrated the same directionality of association between the TF activity and both the TCS feature and image feature. The image validated model list was comprised of 12 triplets, revealing 11 unique DNA loci exerting influence over lymphocyte infiltration through activity perturbation to 8 TFs (Table 1). Notably, 8 out of the top 10 strongest causal models for the TCS phenotype (as ranked by association with TF activity) validated for the lymphocyte density trait, highlighting the excellent predictive potential of our models in the image cohort. The remaining 2 triplets negatively associated with the trait and demonstrated a stronger predictive power in the images.

**Table 1.** CMIF output of genomic drivers and TF perturbations causal for the TCS trait that also predict lymphocyte density in stained tumour sections.

| CNA | TF | TF-TCS Correlation | TF-TCS P value | Model | Type |
| --- | --- | --- | --- | --- | --- |
| RBBP5 | YEATS4 | -0.10 | 0.00 | 1 | TCS |
| CREBBP | ETS1 | 0.62 | 0.00 | 1 | TCS |
| EP300 | ETS1 | 0.62 | 0.00 | 1 | TCS |
| PIAS3 | IRF1 | 0.94 | 0.00 | 1 | TCS |
| PIAS3 | TFEB | 0.80 | 0.00 | 1 | TCS |
| POU2F1 | NR3C1 | 0.60 | 0.00 | 1 | TCS |
| PARP1 | HES1 | -0.85 | 0.00 | 1 | TCS |
| NR5A2 | NFYA | -0.08 | 0.00 | 1 | TCS |
| PATZ1 | BACH2 | 0.81 | 0.00 | 1 | TCS |
| TAL1 | RUNX3 | 0.91 | 0.00 | 1 | TCS |
| CBFB | RUNX3 | 0.91 | 0.00 | 1 | TCS |
| HAX1 | HCLS1 | 0.84 | 0.00 | 1 | TCS |

Of our validated models, notable examples include a process by which *PIAS*3 copy number aberration was found to attenuate the TCS/lymphocyte density through *TFEB* and *I RF*1 activity repression, both of which positively associate with the mentioned traits (Table 1). *CREBBP* and *EP300* were found to exert similar causal pressure on the traits through their action on the TF *ETS1*. This follows from experimental evidence demonstrating that *CREBBP* and *EP*300 form a protein complex *CBP*/*p*300 that is recruited by ETS1 to facilitate its transcription factor functionality [30].

Among the top 8 candidate TFs (Table 1) we identified *N R3C*1: this gene encodes for the glucocorticoid receptor, and influences immune activity through inflammation [31]. This TF was ranked only 95th of 510 in the list of image associations and 27th of 510 in the TCS associations. Its function as a driver of immune infiltration was only elucidated once the causal relationships between genome, signaling and immune phenotypes were modeled together, highlighting the advantage of CMIF over standard pairwise-association approaches.

### Causal model case studies and mechanisms

Our results provide several specific biological examples of causal models of the interaction between cancer signaling and immune activity.

#### EP300 and NCOR1 modulate cytolytic activity through ETS1/SPI1/TP53 network perturbation

Copy number amplification of *EP*300 and *NCOR1* were found to underlie the cytolytic activity trait in both the METABRIC and TCGA cohorts. Interestingly, the original study by Rooney and colleagues [17] found that single nucleotide variants in these genes correlated positively with cytolytic activity in cancer types other than breast. The CMI’s ability to elucidate these mechanisms in breast cancer may be due the higher prevalence of CNA mutations over SNPs in the disease [32]. Furthermore, our method extends the understanding of the association between these DNA-level drivers and the cytolytic score by suggesting they act through perturbations to the activity of *ETS1, SPI*1 and *TP*53.

Our discovery of a positive association between *SPI*1 activity and the CS (*ρ* = 0.7) is consistent with studies demonstrating that *SPI*1 transcribes *CCL*5, a key player in cytolytic activity [33]. Similarly, *ETS*1 deletion in mice has been linked to decreased cytolytic activity in NK cells [34], consistent with our observed positive correlation (*ρ* = 0.58). The association between TP53 activity and cytolysis is not well characterised, although it has been shown that mutant TP53 attenuates cytolytic activity in ovarian and other cancers [17].

The direct correlations between cytolytic activity and EP300 (*ρ* = 0.141) and *NCOR1* (*ρ* = 0.08) are weak, and the additional causal context provided by CMIF was needed to highlight *EP*300 and *NCOR*1 amplification as drivers of cytolytic activity in breast cancer through TF network perturbation.

#### TF drivers of immune localisation regulate adaptive immune pathways

Functional annotation of TF transcriptional targets can elucidate which molecular pathways are over- or underrepresented in the presence of an immune phenotype. We investigated this by partitioning our model set into those where the activity of a TF positively associated with lymphocyte infiltration and those that were negatively associated. We then aggregated the transcription targets of each TF and applied GO term enrichment to functionally annotate the gene sets (Fig. 3A).

**Figure 3.**
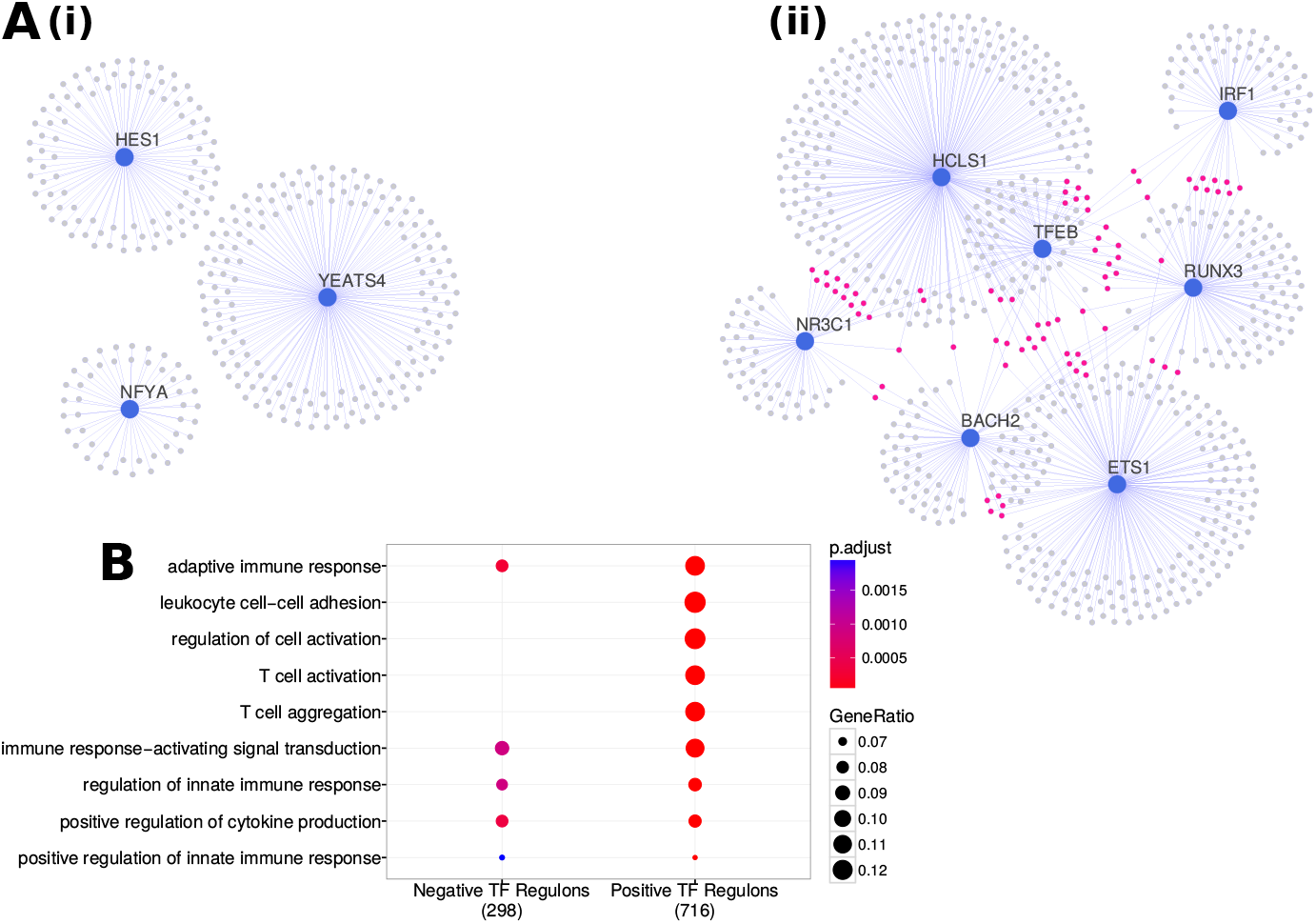
Analysis of TF targets. **A** Network visualisation of the inter-regulon overlap (illustrated through purple dots) between causal TFs that **(i)** down-regulate the TCS/lymphocyte density and **(ii)** those that up-regulate it. Evidently, TFs that causally up-regulate the T cell representation have a greater degree of regulon overlap whereas no intersection is observed for TFs that down-regulate the trait. **B** GO term enrichment analysis highlighting the most significantly annotated terms to the gene sets **A(i)** and **A(ii)** respectively. Whereas regulons pertaining to TFs positively associative with lymphocyte infiltration are more enriched for T-cell related pathways, down-regulators the phenotype are more associated with innate immune system pathways.

For TFs positively associated with lymphocyte recruitment, the term “T cell activation” and “adaptive immune response” (full term list in Additional file 1) were among the top associated pathways (adjusted *P* = 1.7 × 10^−33^ and 2.4 × 10^−28^ respectfully). Interestingly, “antigen processing and presentation” was also ranked highly on the list (adjusted *P* =1.3 × 10^−14^) highlighting the importance of comprehensive antigen recognition in facilitating lymphocyte recruitment. All positively associative TFs in our model space demonstrated target overlap with one another (Fig. 3B(i)). TFs negatively associated with lymphocyte recruitment displayed disjoint sets of target genes (Fig. 3B(ii)) whose aggregate functional annotation was predominantly associated with pathways involved in innate immune cell regulation.

#### Systems driving lymphocyte recruitment stratify by ER status

Differences in magnitude and prognostic relevance of lymphocytic infiltration between ER stratified breast cancer patients have been widely observed [35, 36, 7], but little is known regarding the causal chain of events that gives rise to this discrepancy. We hypothesised that suppressed TF activity in ER+ samples could suggest a possible mechanism by which ER+ tumours evade immune destruction. To investigate this, we used the clinical data for the METABRIC cohort samples to investigate whether genomic drivers of lymphocytic recruitment stratify by ER status.

The copy number profiles of all genomic drivers in our list of models (Table 1) significantly stratified by ER status with the exception of HAX1 (Fig. 4A). For example, genes such as *PIAS*3, *POU2F*1 and *CREBBP* were significantly more amplified in ER+ patients over ER−. These genes downregulate TFs positively associated with lymphocytic activity such as *TFEB*, *IRF*1, *NR3C*1 and *ETS*1 leading to significantly less activity relative to ER− samples (Fig. 4B). Further to this, TAL1 and CBFB were both significantly more amplified in ER− samples over ER+, and subsequently, significantly higher activity observed in the activity of the respective TFs they modulate in ER− samples over ER+. Significantly lower cytolytic activity was also observed in ER+ samples relative to ER− (see Additional file 1), which is unsurprising given that transcriptional targets of TFs regulating the TCS were shown to modulate T cell activation (Fig. 3B).

**Figure 4.**
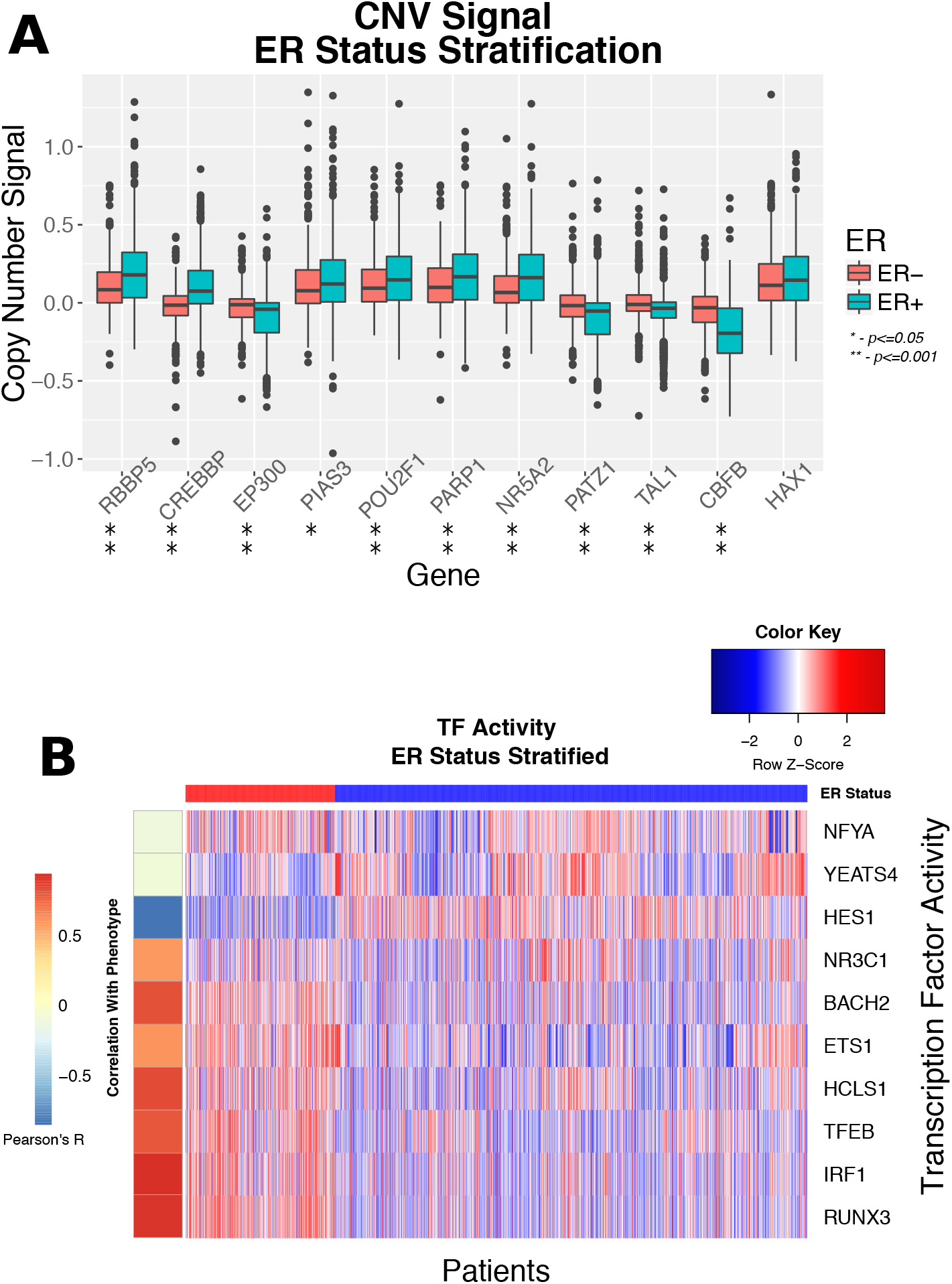
METABRIC ER Stratification of Causal Models. **a** Boxplots highlighting the difference in the normalised DNA copy number signal between ER+/ER− cases in our validated triplet list. Population mean rank difference is computed using the Wilcoxon signed-rank test. The plot shows that 10/11 genes are differentially amplified/deleted between ER+/ER− at *P* ⩽ 0.05. **b** Heatmap highlighting the difference in causal transcription factor activity as stratified by ER status. It can be seen that a large proportion TFs positively associated with lymphocyte infiltration have upregulated activity in the majority of ER− samples. Concurrently, TFs inversely or weakly correlated with the phenotype demonstrate stronger representation in ER+ samples over ER−.

The stratification of these causal events provides compelling evidence of a genomic basis for the ER stratification of lymphocyte infiltration and activity. These results are difficult to infer from association studies alone, highlighting a chief advantage of a deriving causal frameworks from large datasets.

## Discussion & Conclusions

The aim of our study was to dissect interactions between cancer cells and their microenvironment. To achieve this aim, we developed a multistep methodology to inferring directed relationships between signaling activity and immune infiltration in the tumour microenvironment that overcomes limitations of conventional association studies by anchoring the analysis on somatic genomic events.

Our approach uses established methods to estimate TF activity and lymphocyte infiltration (CS) from gene expression data, and proposed a novel score for lymphocytic activity (TCS). Since genes tend to be amplified/deleted together, we use a PPI network as a biological prior to isolate drivers from a list of genes correlated with an imune trait. Causal inference is achieved using a likelihood test, which returns the most likely relationship between the genotype, TF activity and phenotype given the data. Association based methods are widely used in cancer research and our methodology is a step towards a causal and mechanistic understanding of these relationships.

Our analysis consisted of identifying models for the TCS/CS traits in a discovery cohort (METABRIC [11]), validating them in a large independent cohort (TCGA [12]) and further evaluating their predictive utility using orthogonal measures of lymphocyte infiltration from H&E images. The final model list revealed 11 driver genes modulating lymphocyte recruitment into the microenvironment through perturbation to the activity of 8 TFs. Whilst most TFs in our models have been experimentally linked to lymphocyte infiltration, the majority of driver genes we found are novel, highlighting a principal advantage of causal driver discovery over standard association studies. This was further realized with the discovery of *EP*300/*NCOR*1 copy number alterations as drivers of cytolytic activity, whereas SNV mutations in these genes were previously found not to correlate with the trait in breast cancer. Drivers of lymphocytic infiltration were found to stratify by ER status, leading to significant stratification of activity profiles of TFs found to be causal for lymphocyte infiltration. This observation provides evidence supporting a genomic basis for the observed stratification of lymphocyte infiltration and prognostic utility by ER status.

Our approach has several limitations, many of which are technical and relate to the assumptions we had to make for statistical modelling. One such limitation involves measurement errors within the individual data inputs to our integrative analysis. If the margin of error for one variable is wider than that of another, it could potentially lead to the misclassification of the causal-reactive relationship between the two variables. Another limitation arises from the simplicity of the DAG models we design for the interaction between two variables. TFs causal for a trait will regulate genes that are interacting within the context of a much larger network and with feedback controls that need to be accounted for. Although our models successfully predict lymphocyte infiltration in image cohorts, a stronger validation would involve knockdown experiments in mice to to directly observe changes in TF profile and our trait of interest. While our methodology is generally applicable, the details of the statistical model will have to tailored to the specific type of data used in the study, which might not all be normally distributed.

A conceptual limitation is that the whole study is based on one major assumption: genomic events in the cell-autonomous compartment drive the development of the cancer and can thus be used as anchors for causal analysis. This assumption is shared by almost all cancer genomics studies, in particular those that aim to identify genomic drivers of the disease [11, 32]. At the same time, we acknowledge that there could be situations in which microenvironmental changes like inflammation cause genomic events, rather than being caused by them.

Despite these limitations, our method has demonstrated power to recapitulate known mechanisms and has shown results that stay robust when using independent data sets and orthogonal data types. Given our method’s high validation rate between two large and independent datasets and its capacity to predict results supported by the literature, we believe it to be a robust predictor of cancer-immune communication mechanisms in transcriptomic data.

Our analysis was focused on breast cancer, but large efforts like The Cancer Genome Atlas (TCGA) or the International Cancer Genome Consortium (ICGC) provide the same types of data for many other kinds of cancer and thus our methodological framework can be easily be applied to many other forms of the disease. Additionally, the framework is not confined to the CS/TCS metrics as measures of immune activity and can be applied to any other available feature of the microenvironment. Thus, in summary, we have presented an integrative analysis of genomic events, signaling activity and immune markers, which is flexible and can form the foundation for a more mechanistic understanding of tumour-microenvironment interactions across cancer types.

## Materials and Methods

### CMIF Step 1: Data preparation

#### Gene Expression Data

Microarray transcriptomic profiles corresponding to 1980 patients from the METABRIC cohort were downloaded from the European Genome-Phenome archive under the accession id: EGAD00010000268 (https://ega-archive.org/datasets/EGAD00010000268). The issue of multiple probes mapping to the same gene was addressed by selecting the probe with the highest variance. RNA-seq count data comprising 1154 BRCA samples was downloaded from the TCGA archive (https://tcga-data.nci.nih.gov/tcga) and processed using a two-step process: applying the variance stabilising transform and quantile normalising the matrix with respect to the METABRIC gene expression distribution. This was done to correct for the large heteroscedasticity between genes and make the expression distributions more comparable.

#### DNA copy number aberrations

METABRIC Affymetrix SNP 6.0 data were downloaded from the same resource as the transcriptomic data. SNP array genomic positions were mapped to gene symbols using the hg18 build. TCGA GISTIC2 gene-level, zero-centered, focal copy number calls for each patient were accessed from GDAC Firehose (http://gdac.broadinstitute.org/).

#### Transcripton Factor Network Enrichment

To infer TF activity, we used a coexpression network [37] for 788 experimentally verified TFs derived using mutual information and used this to calculate the activity of each TF for each sample using the R package viper (virtual inference of protein activity by enriched regulon analysis) [16] using default parameters. VIPER tests the activity of each TF by examining the relative transcript abundance of known targets genes, collectively referred to as a “regulon”. An activity score for each TF is computed using an enrichment analysis method that includes a probabilistic weighting using TF-gene association likelihoods.

#### Immune trait inference from transcriptomic data

To gauge the degree of T cell infiltration in transcriptomic data, we introduce the T cell score (TCS) as the geometric mean of *CD3D, CD4, CD8A* and *CD8B* expression. In addition to being well established T cell markers, they are expressed primarily in cells of a haematopoietic lineage with little noise contamination from the rest of the tumour. To measure cytolytic activity from transcriptomic data, we define the cytolytic score (CS) as the geometric mean of *GZMA* and *PRF*1 as per wok done by Rooney and colleagues [17]. In both cases, the geometric mean is chosen over the arithmetic mean given its reduced sensitivity to outliers.

#### Protein-protein interactions

A network detailing protein-protein interactions was downloaded from the STRINGdb resource (http://string-db.org/) and ENSEMBL identifiers were mapped to HUGO gene symbols using the R biomaRt package.

### CMIF Step 2: Triplet Initialisation

Here we describe our approach for deriving pairwise associations between CNA profiles, TF activity and immune phenotypes. These triplet skeletal graphs are the input to the model scoring step of the CMIF.

#### Significance of assocations

All associations in this manuscript are computed using the Pearson correlation coefficient and *p*-values are calculated using Student’s test unless explicitly stated otherwise.

#### Computing undirected triplet graphs

Undirected skeletal graphs were constructed by computing pairwise associations between the CNA profiles, TF activity and the phenotype of interest. To reduce the number of hypotheses to test, we only considered pairs of TFs and CNAs, if the corresponding proteins showed protein-protein interaction.

METABRIC was taken as our discovery cohort and thus association *P*-values were adjusted using the Benjamini-Hochberg (BH) procedure [21] and a conservative significance threshold defined at *P* ⩽ 10^−3^. Findings were said to be validated if reproduced at unadjusted *P* ⩽ 0.05 in the TCGA cohort.

Skeletal graphs between triplets of variables were passed to the model scoring step if all three pairwise associations in the graph demonstrated significance.

### CMIF Step 3: Model scoring

#### Likelihood Function Definitions

Our approach uses Bayesian networks and likelihood models to determine which relationship between the variables is best supported by the data. Assuming the conditional probability distribution of the future state of the system depends only on the present state, we can write our joint probability distribution of models *M*_1_, *M*_2_ and *M*_3_ as

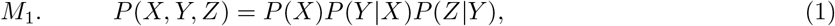

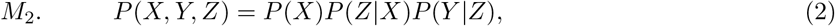

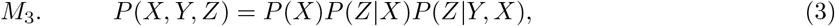

where X, Y and Z correspond to the copy number profile, transcription factor activity and immune phenotype measurements respectively. We assume that Y and Z are normally distributed with a constant variance such that the likelihood of each model can be described by multivariate Gaussian density functions. Individual component likelihoods are summed over all copy number states (*J* = {*Amplified, Deleted, Neutral*}) and the likelihood of each joint distribution given the parameterisation is computed by multiplying over the entire sample space *N* as such:

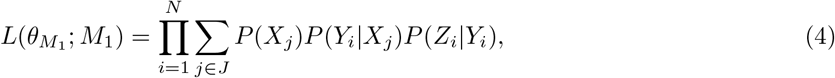

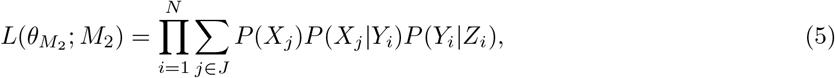

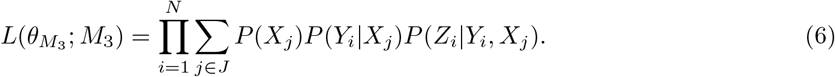

Individual component definitions are described in the Additional file 1.

#### Maximum Likelihood Estimation and Model Selection

The joint distribution likelihoods are maximised over the parameter space using the maximum likelihood estimation algorithm as implemented by the *optim* function in R. We then compute the Akaike information criterion (AIC) as

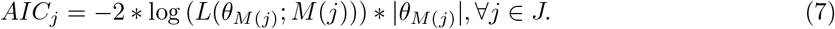

The model with the smallest AIC is the strongest causal candidate as given by the data.

### Validation Approaches

The robustness of our analysis is evaluated with several validation strategies that use orthogonal datasets such as H&E images and flow cytometry data.

#### T Cell Score Validation using Flow Cytometry

Gene expression profiles for 20 peripheral blood mononuclear cell admixture samples and their corresponding flow cytometry profiles as measured by Newman *et al* [9] were downloaded from (http://cibersort.stanford.edu). Pearson’s test was then used to infer correlations between flow cytometry measurements for 9 subsets of leukocytes and the TCS measurements for all 20 samples.

#### H&E Section Validation

We made use of the image dataset published by Yuan and colleagues [13], comprising the segmented H&E stained primary tissue sections of 564 patients sampled from the METABRIC cohort. Segmented objects were classified using a SVM trained by an expert pathologist and metrics pertaining to the absolute number of lymphocytes and the lymphocyte density relative to the number of overall objects were measured. The absolute number of lymphocytes were then log-transformed to enable more robust comparison across the patient space. Furthermore, this transformation ensures input to the MLE process exhibits a similar range of values and ensure that the algorithm can be initialised with the same parameters for all cohorts. Finally, CMIF was run with the lymphocyte statistics, their paired copy number profiles and TF activities.

### Additional analyses

#### GO Term Enrichment Analysis

GO term enrichment analysis is used to characterise groups of TFs causal for our observed phenotypes. Causal TF regulators of immune infiltration were split into two groups given by the directionality of their association with the TCS phenotype. Their regulons were aggregated and GO term enrichment was performed using the *clusterProfiler* package in R [38]. Further details can be found in the Supplementary Statistical Analysis (Additional file 1).

### Code & Data Availability

The R script [14] implementation for the CMIF is provided in the Supplementary R Sweave file (Additional file 1). Additional file 1 can also be used to reproduce all figures and results of the described case studies.

Data accession details are provided in the Materials and Methods section. Furthermore, the datasets used can be downloaded directly using either code available in the R Sweave file (Additional file 1) or from the github repository https://github.com/databro/Chlon2017.

### Ethics approval

Ethics approval was not needed for this study.

## Acknowledgements

We thank Dr. Federico Giorgi (University of Cambridge) for fruitful discussions and project reproducibility testing.

## Competing interests

The authors declare that they have no competing interests.

## Author’s contributions

LC contributed to project conceptualisation, data curation, formal analysis, investigation, methodology, validation and visualisation. LC & FM contributed to writing, review and editing. FM contributed to funding acquisition, project administration and supervision.

## Funding

This work was funded by the Cancer Research UK and Engineering and Physical Sciences Research Council Imaging Centre in Cambridge and Manchester grant (C197/A16465).

## Additional Files

Additional file 1

Zipped file containing a set of R code, results, and datasets to fully reproduce the results and analysis outlined in this manuscript.

